# Versican controlled by Lmx1b regulates hyaluronate density and hydration for semicircular canal morphogenesis

**DOI:** 10.1101/2024.05.07.592968

**Authors:** Yusuke Mori, Sierra Smith, Jiacheng Wang, Akankshi Munjal

## Abstract

During inner ear semicircular canal morphogenesis in zebrafish, patterned canal-genesis zones express genes for extracellular matrix component synthesis. These include hyaluronan and the hyaluronan-binding chondroitin sulfate proteoglycan Versican, which are abundant in the matrices of many developing organs. Charged hyaluronate polymers play a key role in canal morphogenesis through osmotic swelling. However, the developmental factor(s) that control the synthesis of the matrix components and regulation of hyaluronate density and swelling are unknown. Here, we identify the transcription factor, Lmx1b, as a positive transcriptional regulator of hyaluronan, Versican, and chondroitin synthesis genes crucial for canal morphogenesis. We show that Versican regulates hyaluronan density through its protein core, whereas the charged chondroitin side chains contribute to the osmotic swelling of hyaluronate. Versican-tuned properties of hyaluronate matrices may be a broadly used mechanism in morphogenesis with important implications for understanding diseases where these matrices are impaired, and for hydrogel engineering for tissue regeneration.

**Summary Statement:** Here, we reveal the function of the hyaluronan-binding proteoglycan, Versican, and its chondroitin sulfate side chains in tuning the density and hydration of the hyaluronate-rich ECM to generate force, under the control of the transcription factor Lmx1b for successful inner ear semicircular canal morphogenesis in zebrafish.

## Introduction

Recent studies on developing tissues across species have demonstrated the contributions of extracellular matrices (ECM) in driving shape changes during morphogenesis. These studies revealed that the protein and glycoprotein components, such as collagens, laminins, and fibronectins, form fibrous networks in the ECM, providing a stiff substrate for remodeling tissues (Diaz-de-la-Loza and Stramer, 2024; Wu et al., 2023). Glycosaminoglycans (GAG) and proteoglycans are equally abundant viscoelastic ECM components, but their functions are less appreciated, even though they can be active drivers for tissue remodeling due to their physical properties(Richter et al., 2018; Toole, 1981). A striking example is the GAG hyaluronic acid (HA), which powers the morphogenesis of semicircular canals in the inner ear(Haddon and Lewis, 1991; Munjal et al., 2021). The genes required for HA production, *UDP-glucose-6-dehydrgenase* (*ugdh*) for substrate precursor synthesis and *HA synthase 3* (*has3)* for substrate polymerization, are expressed locally in the semicircular canal genesis zones of the embryonic inner ear during morphogenesis(Munjal et al., 2021). Has3 resides in the plasma membrane and extrudes large negatively charged hyaluronate polymers that osmotically swell to remodel the overlying cells into buds. Tissue morphogenesis driven by the local swelling of hyaluronate-rich ECMs may be a conserved mechanism in other organs in which they are present, such as the heart, eye, and neural tube (Michaut et al., 2022; Nakamura and Manasek, 1981; Toole, 2001; Vignes et al., 2022). In addition, hyaluronate-based gel matrices are extensively studied materials for biomedical applications such as joint lubrication and injectable implants for cosmetic treatments(Yasin et al., 2022). Yet, how the functions and activities of hyaluronate-based ECMs are regulated during morphogenesis *in vivo* remains to be fully investigated.

The primary structure of HA polymers is simple, but the physical properties of the macromolecule can be regulated by binding partners, leading it to acquire complex secondary structures(Cowman et al., 2015; Richter et al., 2018). One such binding partner is the chondroitin sulfate proteoglycan Versican, which non-covalently interacts with HA to form large aggregates(LeBaron et al., 1992). Many developing tissues with a hyaluronate-rich ECM, such as the heart(Peal et al., 2009; Walsh and Stainier, 2001), eye(Kwan, 2014), and inner ear(Geng et al., 2013), also express Versican. Evidence shows that Versican plays a role in heart morphogenesis since embryos lacking Versican die with cardiac defects (Mjaatvedt et al., 1998; Yamamura et al., 1997). These functions are mediated by Versican’s interaction with HA(Derrick and Noel, 2021; Islam and Watanabe, 2020; Wight, 2017). However, the mechanisms underlying the functions of Versican and its chondroitin sulfate side chains in the morphogenesis of other organs still need to be elucidated. Moreover, the developmental factors(s) that upregulate the synthesis of the HA and Versican-rich ECM during semicircular canal morphogenesis have remained unknown.

The inner ears of all jawed vertebrates, including humans and zebrafish, have three mutually orthogonal semicircular canals that sense balance and angular acceleration(Groves and Fekete, 2012). Canal development is understudied due to the inaccessibility of the embryonic inner ear in most model organisms. Zebrafish provide an unprecedented opportunity to study inner ear development(Baxendale and Whitfield, 2016). Their embryos are optically transparent, and the tissue that gives rise to the adult inner ear, the otic vesicle (OV), is not obstructed by the presence of middle and outer ears, rendering it accessible to live or fixed whole-mount imaging. The OV is a small, single-layered epithelium enclosing an endolymph-filled lumen(Whitfield, 2015). Semicircular canal morphogenesis begins two days post fertilization when six canal-genesis zones, which are the *ugdh and has3* expressing regions in the non-sensory dorsal OV, sequentially bud into the lumen(Munjal et al., 2021). Adjacent bud pairs fuse to form pillars, which serve as hubs of the canals(Alsina and Whitfield, 2017). Three sensory cristae sit at the base of each canal to detect endolymph flow during head rotations (angular acceleration).

It is known that LIM homeobox-containing transcription factors, such as Lmx1a and Lmx1b, acting through antagonistic interactions with Notch signaling, pattern the non-sensory (canal-forming) and sensory (cristae-forming) domains of the mouse inner ear (Abello et al., 2007; Brown and Groves, 2020; Chizhikov et al., 2021; Mann et al., 2017; Nichols et al., 2008) and the zebrafish OV expresses two paralogs, *lmx1ba* and *lmx1bb*. Evidence suggests a role for Lmx1b in regulating ECM synthesis during canal morphogenesis. These include ChIPseq data for *Has3* and *Vcan* from mouse limb buds (Haro et al., 2017) and the fact that a loss-of-function mutant of *lmx1bb* has abnormally shaped buds during zebrafish canal development(Obholzer et al., 2012; Schibler and Malicki, 2007; Swinburne et al., 2018).

Here, we explore the role of Versican and Lmx1b in the morphogenesis of the semicircular canals of zebrafish embryos. We demonstrate that Lmx1b patterns the expression of HA and Versican synthesis genes, thereby playing a crucial role in this process. Additionally, we uncover the essential functions of Versican in hyaluronate-powered tissue remodeling. The protein core of Versican, expressed by paralogs, *vcana,* and *vcanb,* is necessary to accumulate HA in the ECM and can tune hyaluronate-ECM density. By contrast, the covalently attached charged chondroitin sulfate (CS) side chains of Versican, polymerized by the enzyme *chsy1,* contribute to the osmotic swelling and hydration of the hyaluronate-Versican aggregates, thereby regulating the pressure buildup in this ECM.

## Results

### *Lmx1ba* and *lmx1bb* are differentially patterned in the non-sensory domain and necessary for semicircular canal morphogenesis

Previous studies have reported expression of the two paralogs of Lmx1b: *lmx1ba* (previously *lmx1b.2)* and *lmx1bb* (previously *lmx1b.1)* in the zebrafish OV using low-resolution *in situ hybridization*(Burzynski et al., 2013). We, therefore, studied their expression dynamics in more detail, using multiplexed hybridization chain reaction (HCR) fluorescent *in situ* hybridization (FISH)(Choi et al., 2018), a quantitative technique, at critical stages of canal development: 40 hpf (pre-budding), 50 hpf (anterolateral, posterolateral, anterior and posterior budding), 60 hpf (fusion of anterolateral and anterior buds, and posterolateral and posterior bud extension), 65 hpf (anterior and posterior pillar formation and ventral budding), and 72 hpf (anterior, posterior, and ventral pillar formation) **(Figure 1A)**. As a positive control, we use *col2a1a,* expressed in most cells in the OV(Munjal et al., 2021) **(Figure 1A)**. Strikingly, *lmx1ba* and *lmx1bb* are expressed in the OV before morphogenesis begins. While *lmx1bb* is more broadly expressed, *lmx1ba* transcripts are restricted to the canal-genesis zones of the lateral, anterior, posterior, and ventrolateral regions in the dorsal OV **(Figure 1A)**. The expression of *lmx1ba* is lost after bud fusion, while that of *lmx1bb* diminishes but continues after bud fusion and pillar formation **(Figure 1A)**. These data show that *lmx1bb* and *lmx1ba* are differentially patterned during secircular canal morphogenesis.

**Figure 1:**
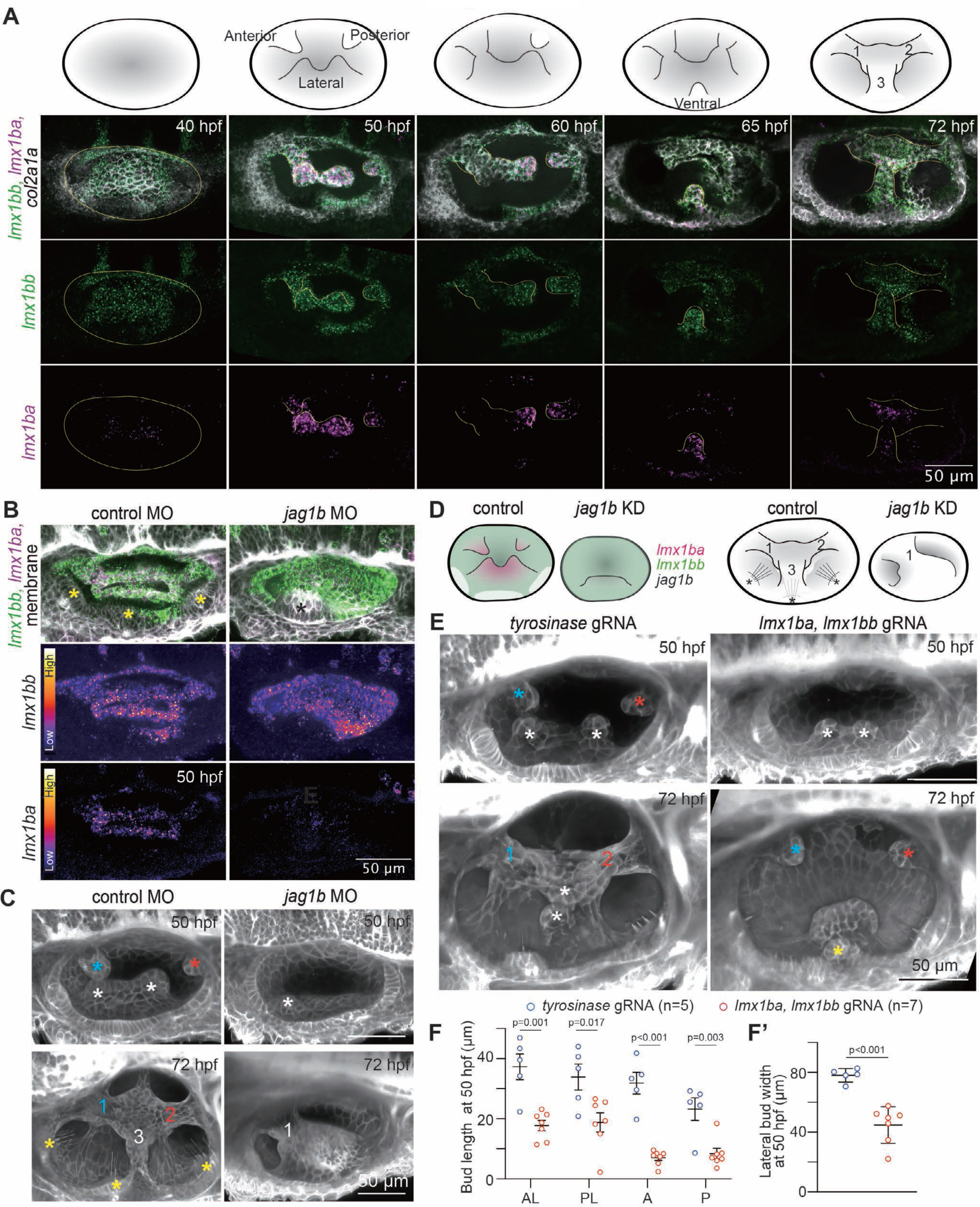
Genes encoding the transcription factor Lmx1b, *lmx1ba, and lmx1bb,* are differentially patterned and necessary for semicircular canal morphogenesis. **A)** Maximum intensity projections of OVs at select time points stained with multiplex *in situ* probes against *lmx1bb* (green), *lmx1ba* (magenta), and *col2a1a* (white). The z-volume and the contrast of each time point are individually set to capture all buds and get better visualization. Illustrations show typical morphological features of the OVs at relevant stages of semicircular canal morphogenesis. Yellow lines indicate buds or pillar. Scale bar, 50 μm. **B)** Maximum intensity projections of OVs of membrane-NeonGreen-expressing embryos injected with control or *jag1b* MO at 50 hpf stained with multiplex in situ probes against *lmx1bb* (green) and *lmx1ba* (magenta). The z-volume of each condition is individually set to capture all buds and show the OVs morphology. Heatmaps represent fluorescent intensity with the same contrast for each probe of embryos injected with control and *jag1b* MO. Crista and macula are marked by yellow and black asterisks, respectively. Scale bars, 50 μm. **C)** 3D-rendered OVs at 50 or 72 hpf from membrane-NeonGreen-expressing embryos injected with control or *jag1b* MO. Scale bar, 50 μm. Anterior, posterior, and lateral buds are marked by blue, red, white, and asterisks, respectively. Yellow asterisks mark hair cells. Scale bar, 50 μm. **D)** Illustrations showing the difference in gene expression of *lmx1ba* and *lmx1bb* at 50 hpf (left) or SCC morphology at 72 hpf (right) between embryos injected with control and *jag1b* MO. Asterisks mark cristae. **E)** 3D-rendered OVs at 50 or 72 hpf from membrane-mCherry-expressing embryos injected with *tyrosinase* or *lmx1b-*double gRNA. Anterior, posterior, lateral, and ventral buds are marked by blue, red, white, and yellow asterisks, respectively. Scale bar, 50 μm. **F and F’)** Quantification of anterior bud length (F) and lateral bud width (F’) in embryos injected with *tyrosinase* or *lmx1b*-double gRNA at 50 hpf. Data are mean ± SE. In the absence of buds, bud lengths correspond to cell lengths. “n” denotes the number of buds from individual embryos measured per condition. p values as labeled (unpaired two-tailed Student’s t-test).

The three semicircular canals form from the non-sensory domain of the OV, while the three cristae at the base of each canal form from the sensory domain(Higuchi et al., 2019; Whitfield, 2015). Given the known functions of the sensory domain in patterning the non-sensory domain(Abello et al., 2007; Brown and Groves, 2020; Mann et al., 2017), we hypothesized that the cristae are required to pattern *lmx1ba* and *lmx1bb* gene expression. The sensory cristae express neither *lmx1ba* nor *lmx1bb* (**Supp Figure 1)**. They do, however, express *jagged1b (jag1b)* encoding a ligand for Notch signaling, as previously reported(Obholzer et al., 2012) (**Supp Figure 1**). Morpholino-mediated *jag1b* knockdown successfully inhibited cristae formation as expected and impaired canal morphogenesis, as previously reported(Ma and Zhang, 2015)**(Figures 1B-D)**. Significantly, *jag1b* knockdown resulted in a broader expression of *lmx1bb*, likely due to the absence of cristae **(Figure 1B)**. Conversely, *jag1b* knockdown resulted in reduced expression of *lmx1ba* **(Figure 1B)**. These data indicate that the differential patterning of *lmx1ba* and *lmx1bb* in the canal genesis zone of the OV is under Notch signaling control (**Figure 1D**). Next, we tested their role in canal morphogenesis.

A null mutant of *lmx1bb, jj410,* with a premature stop codon, has been reported to have abnormal bud formation (Obholzer et al., 2012). We therefore tested the contributions of *lmx1ba* and *lmx1bb* separately and together on bud formation using CRISPR-Cas9 gene editing in F0 embryos. We injected a pool of three single guide RNAs (gRNAs) against each paralog (**Supp Figure 2A**). As a control, we used a gRNA against *tyrosinase*(Sorlien et al., 2018), resulting in pigmentation loss in 90% of the injected embryos (**Supp Figure 2B**). A heteroduplex mobility assay confirmed successful gene-editing by all gRNAs (**Supp Figure 2C**). At 72 hpf, 10% of *lmx1ba* and 37% of *lmx1bb* gRNA injected F0 embryos showed abnormal canal development (**Supp Figure 2D**), while co-injection of gRNAs against *lmx1ba* and *lmx1bb* resulted in 69% of embryos with abnormal canal development. RT-PCR confirmed a 52% and 33% reduction of *lmx1ba* and *lmx1bb*, respectively, in the double knockdown embryos (**Supp Figure 2E**). These data suggest that the two paralogs compensate for each other’s functions. Therefore, we deployed the double knockdown strategy hereafter to investigate the combined loss-of-functions of *lmx1ba* and *lmx1bb* in a single generation. Canal morphogenesis was severely inhibited in the *lmx1ba* and *lmx1bb* double knockdown embryos: the lateral buds formed with a delay and were significantly smaller in length and width, and the anterior and posterior buds in most embryos were either absent or significantly smaller (**Figures 1E and 1F**). These embryos had no pillars at 72 hpf compared to the three pillars in control embryos (**Figure 1E**). Altogether, these data show that *lmx1ba* and *lmx1bb* are required for the morphogenesis of the semicircular canals.

### *Lmx1ba* and *lmx1bb* regulate HA and Versican synthesis genes for morphogenesis

HA is a crucial driver for budding morphogenesis during canal development(Haddon and Lewis, 1991; Munjal et al., 2021). We therefore asked if the failure of budding morphogenesis in the *lmx1ba* and *lmx1bb* double knockdown condition is due to the role of the transcription factors in patterning the expression of the HA synthesis gene *has3,* in the canal genesis zones. In wildtype OVs, *has3* expression increases during the bud extension stage and decreases during bud fusion(Munjal et al., 2021), corresponding to ∼50- and ∼60 hpf for the anterior bud, respectively. We therefore compared *has3* gene expression levels in the anterior canal genesis zone at 50 hpf between *tyrosinase* control and *lmx1bb* and *lmx1ba* double knockdown embryos (**Figure 2B**). We first validated successful gene editing in individual embryos using a T7 endonuclease assay (**Figure 2A**). The HCR-FISH protocol for control and validated double knockdown embryos was carried out in the same tube to minimize variability and facilitate a quantitative gene expression comparison (**Figure 2A**). The anterior bud showed a 44% reduction in *has3* gene expression of *lmx1ba* and *lmx1bb* double knockdown embryos compared to controls (**Figures 2B and 2C**). Qualitatively, some cells in the lateral canal genesis zone continued to express *has3*, although the size of the lateral bud was significantly smaller than controls (**Figures 1E-F, and Supp Figure 3D**). Together, these data show a requirement for Lmx1b in budding morphogenesis at least in part via the expression of *has3*.

**Figure 2:**
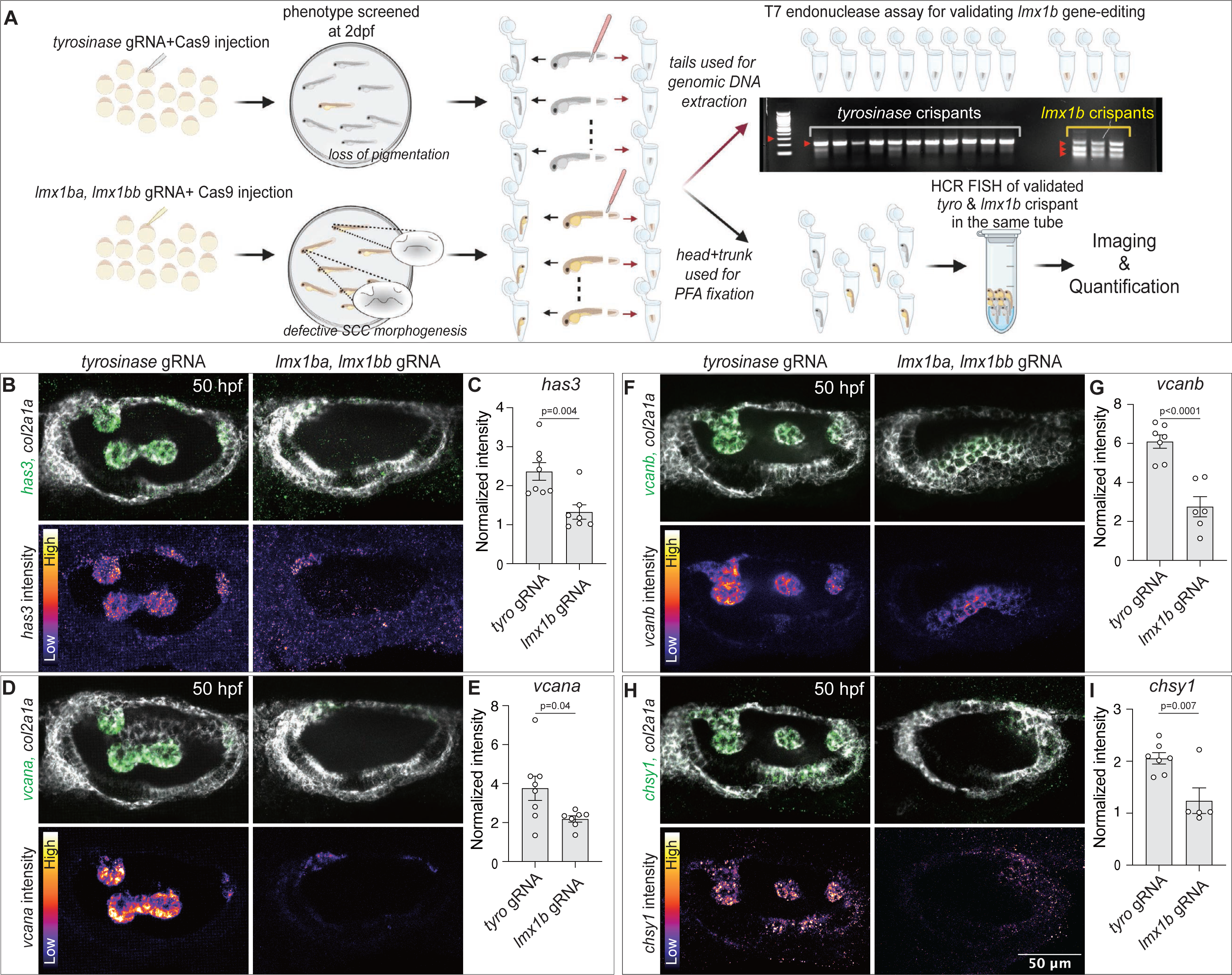
*Lmx1ba* and *lmx1bb* regulate HA and Versican synthesis genes for morphogenesis. **A)** The workflow illustrates screening and validation of F0 gRNA/Cas9-injected embryos, followed by a multiplex in situ hybridization experiment. gRNA/Cas9-injected embryos were screened at 50 hpf using their inner ear morphology. A small piece from the tail was clipped to assess genome editing using a T7 endonuclease assay. The remaining embryo body was fixed with PFA. Individually validated embryos from controls and *lm1ba* and *lmx1bb* knockdown were pooled in a single tube to perform HCR-FISH. **(B-I)** Effect of *lmx1b*-double knockdown on ECM-associated gene expression in OV. Maximum intensity projections of OVs and quantification of fluorescent probe intensity of *has3* (B and C), *vcana* (D and E), *vcanb* (F and G), and *chsy1* (H and I) probes. *col2a1a* probe and the other ECM-associated gene probes are shown as white and green, respectively. The z-volume of each condition is individually set to capture all buds and show the OVs morphology. Heatmaps represent fluorescent intensity with the same contrast for each probe of embryos injected with *tyrosinase* or *lmx1b*-double gRNA. Scale bar, 50 μm. Data are mean ± SE. “n” denotes the number of buds from individual embryos measured per condition. p values as labeled (unpaired two-tailed Student’s t-test).

Lmx1b is best studied for its role in limb dorsalization as a transcription factor that broadly regulates ECM synthesis, including proteoglycans like Versican (Feenstra et al., 2012; Haro et al., 2017). We, therefore, asked whether Versican in the OV is also regulated by *lmx1ba* and *lmx1bb*. The protein core of the zebrafish Versican is encoded by two paralogs, *vcana* and *vcanb*(Geng et al., 2013). The synthesis of CS side chains requires *ugdh* for substrate precursor and several glycosyl transferases, including *chondroitin synthase 1* (*chsy1),* for glycosylation and substrate polymerization(Filipek-Gorniok et al., 2013; Li et al., 2010). Consistent with previous literature(Filipek-Gorniok et al., 2013; Geng et al., 2013), and like *has3* (Munjal et al., 2021), *vcana, vcanb,* and *chsy1* are expressed exclusively and transiently at the canal genesis zones in control embryos (**Supp Figure 3A-C**). Moreover, the expression of all three genes can be detected before canal morphogenesis begins (**Supp Figure 3A**). Strikingly, *lmx1ba* and *lmx1bb* double knockdown embryos exhibited 42% reduction in *vcana*, 55% reduction in *vcanb*, and 40% reduction in *chsy1* expression at the anterior canal genesis zone (**Figures 2D-I**). In addition, as seen for the *has3* knockdown, some cells in the lateral canal genesis zone of the double knockdown embryos continued to express *vcana, vcanb,* and *chsy1;* this continuation was associated with smaller bud sizes compared to controls (**Figures 1E-F and Supp Figure 3D**). These data show that Lmx1b is necessary to express genes driving HA and Versican synthesis during semicircular canal morphogenesis. However, the functions of Versican during canal morphogenesis have remained unknown, and we therefore test this next.

### Versican is necessary for semicircular canal morphogenesis

The interactions of Versican with HA are critical for its function in heart morphogenesis in zebrafish, medaka, and mice (Derrick and Noel, 2021; Islam and Watanabe, 2020; Nandadasa et al., 2014). To address the role of Versican in canal morphogenesis, we first carried out CRISPR-Cas9-or morpholino (MO)-mediated knockdown of the protein core encoded by *vcana* and *vcanb* (**Supp Figure 4A**). Using heteroduplex mobility assay and RT-PCR, we validated successful gene editing and significant mRNA reduction by CRISPR-Cas9 in F0 embryos (**Supp Figures 4B-C**). Separate knockdown of *vcana* and *vcanb* with CRISPR-Cas9 did not perturb development (**Supp Figures 4D**). Similarly, single knockdowns of *vcana* and *vcanb* with MO only showed a mild defect in which canal morphogenesis continued with a delay (**Supp Figures 4E**). These results suggest a compensation between the two paralogs. We, therefore, simultaneously knocked down *vcana* and *vcanb* by either using double MO or double guide RNAs against both paralogs. Semicircular canal morphogenesis was severely affected in the *vcana* and *vcanb* double morphants or crispants, with smaller buds at 50 hpf and impaired morphogenesis with no pillar formation at 72hpf (**Figures 3A-D**).

**Figure 3.**
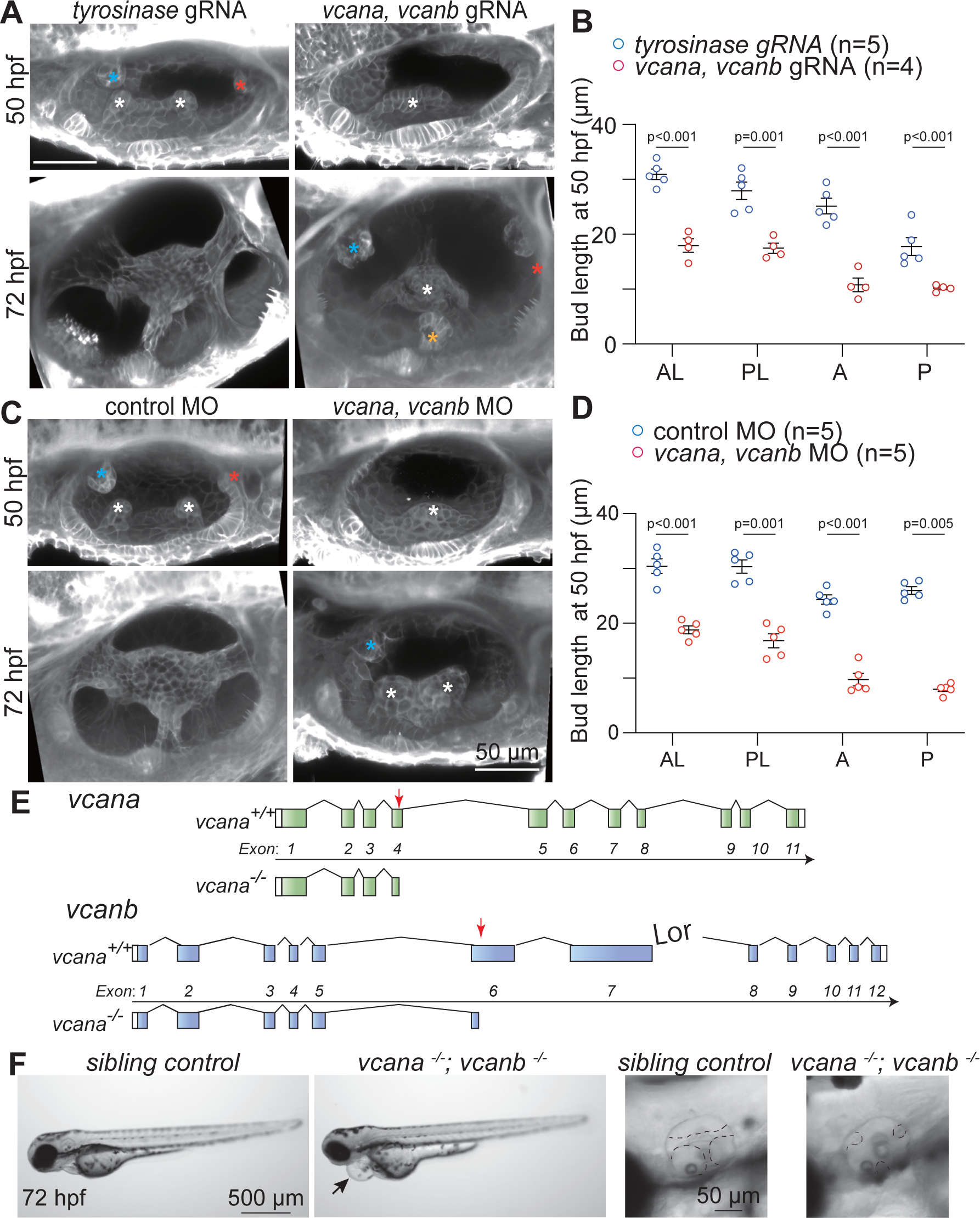
Versican, a chondroitin sulfate proteoglycan, is required for semicircular canal morphogenesis. **A and C)** 3D-rendered OVs at 50 or 72 hpf from membrane-NeonGreen-expressing embryos injected with *tyrosinase* (left) or *versican*-double (right) gRNA (**A**), and control (left) or *versican*-double MO (right) (**C**). Anterior, posterior, lateral, and ventral buds are marked by blue, red, white, and yellow asterisks, respectively. Scale bar, 50 μm. **B and D)** Quantification of anterior bud length in embryos injected with *tyrosinase* or *versican*-double gRNA (**B**), and control or *versican*-double MO **D**) at 50 hpf. Data are mean ± SE. In the absence of buds, bud lengths correspond to cell lengths. “n” denotes the number of buds from individual embryos measured per condition. p values as labeled (unpaired two-tailed Student’s t-test). **E)** Target sites for sgRNAs at the *vcana* or *vcanb* locus used to generate stable *versican*-double KO mutants. **F)** Bright-field images of sibling control and *versican-*double-KO mutant embryos at 72 hpf obtained from *vcan^-/+^; vcanb^-/-^parents. The arrow indicates epicardial edema in the versican*-double KO mutant embryo. The dashed lines in the magnified images indicate the outlines of pillars or buds in the OV. Scale bar, 50 or 500 μm.

**Figure 4.**
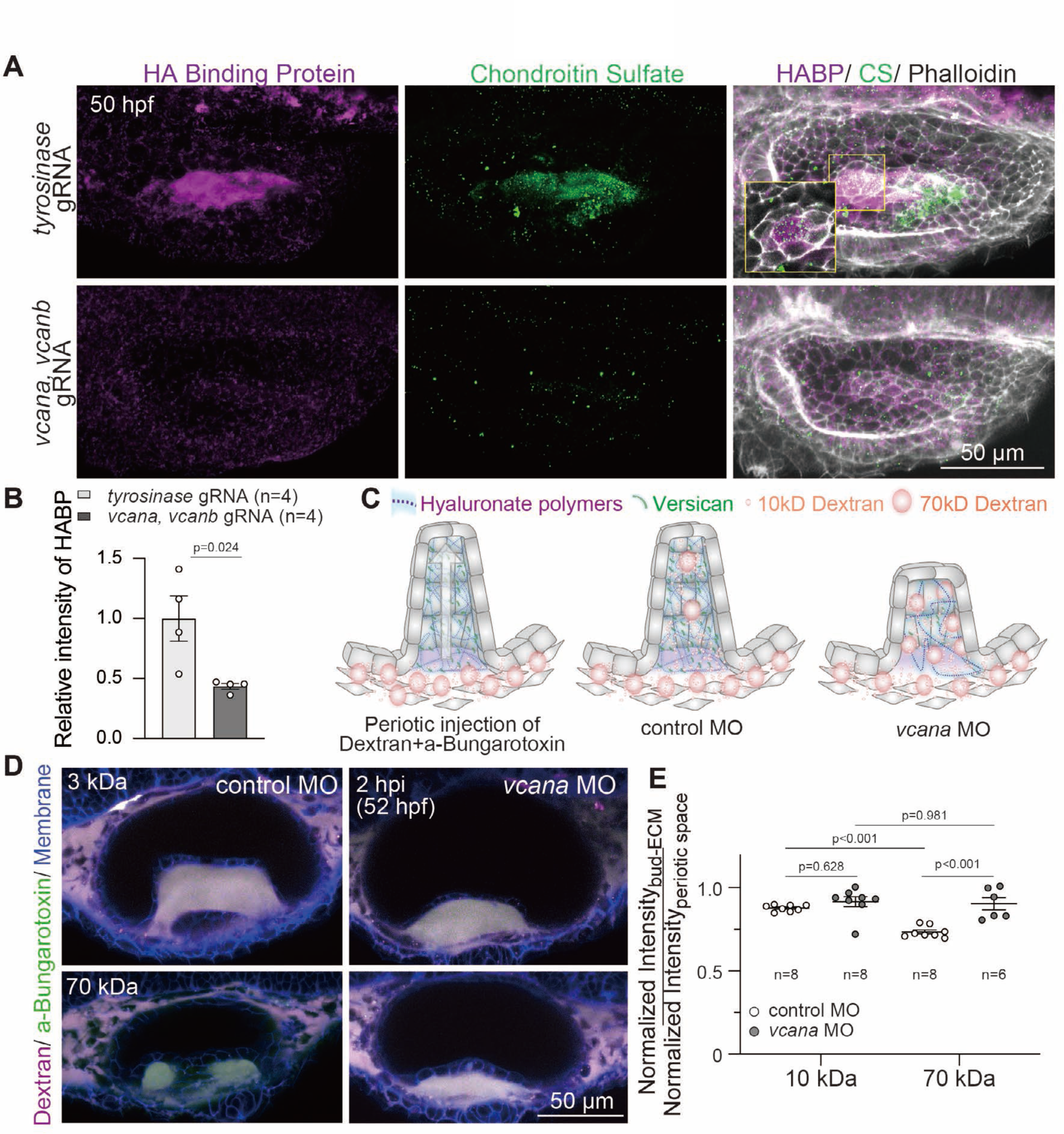
Versican regulates density of the bud-ECM via accumulation of HA. **A-B)** 3D-rendered OVs showing hyaluronic acid (HA), chondroitin sulfate, and F-actin using HA-binding protein, anti-chondroitin sulfate antibody, and phalloidin, respectively, in embryos injected with *tyrosinase* or *versican*-double gRNA at 50 hpf. Insets show 2D sections of the antero-lateral buds. Scale bar, 50 μm. **B)** Quantification of fluorescent intensity of HABP in embryos injected with *tyrosinase* or *versican*-double gRNA. 5 μm-thickness maximum intensity projection images of HABP in the lateral bud region were used for the quantification. Data are mean ± SE. “n” denotes the number of embryos measured per condition. p values as labeled (unpaired two-tailed Student’s t-test). **C)** Illustration showing different sizes of dextran injected in the periotic region percolating into the bud-ECM region in control and *vcana* knockdown conditions. **D)** Composite of 2D sections of OVs showing percolation of dextran from periotic region to into bud-ECM region at 2 hours post-injection (hpi). Different sizes of Texas-red dextran 3 kDa or 70 kDa (magenta) with approximate Stokes radii 1 or 5 nm, respectively, were co-injected with aBt (green) in *membrane-NeonGreen (blue)-expressing control or vcana MO-injected* embryos. aBt colocalizes with all three dextran sizes in the periotic region (white). Contrast is the same across embryos. Scale bar, 50 μm. **E)** Quantification of fluorescent intensities of different sizes of dextran in the bud-ECM region normalized to their intensities in the periotic space. “n” denotes the number of embryos measured per condition. p values as labeled (one-way ANOVA with Tukey’s test).

We also generated stable double mutants of *vcana* and *vcanb* using CRISPR-Cas9 (**Figure 3E and Supp Figure 5A**). Guide RNAs targeting exon 4 in *vcana* or exon 6 in *vcanb* were co-injected, generating seven bp deletions resulting in frameshifts that induced premature stop codons (**Supp Figures 5B-C**). F2 fish with genotypes vcana*+/-* and *vcanb-/-* were incrossed to obtain F3 fish with double knockout genotype (*vcana -/-, vcanb-/-*). At 72 hpf these F3 double mutants showed incomplete semicircular canal morphogenesis like F0 *vcana* and *vcanb* double knockdown embryos (**Figure 3F**). We named the double mutant *memai*, Japanese for dizzy– a common symptom of semicircular canal disorders. The siblings of *memai* with genotypes *vcana*+/+ or *vcana*+/- did not show canal morphogenesis defects (**Figure 3F and Supp Figure 5D**), corroborating the functional redundancy of the two paralogs during canal morphogenesis. These data show that the protein core of Versican is necessary for canal morphogenesis.

**Figure 5.**
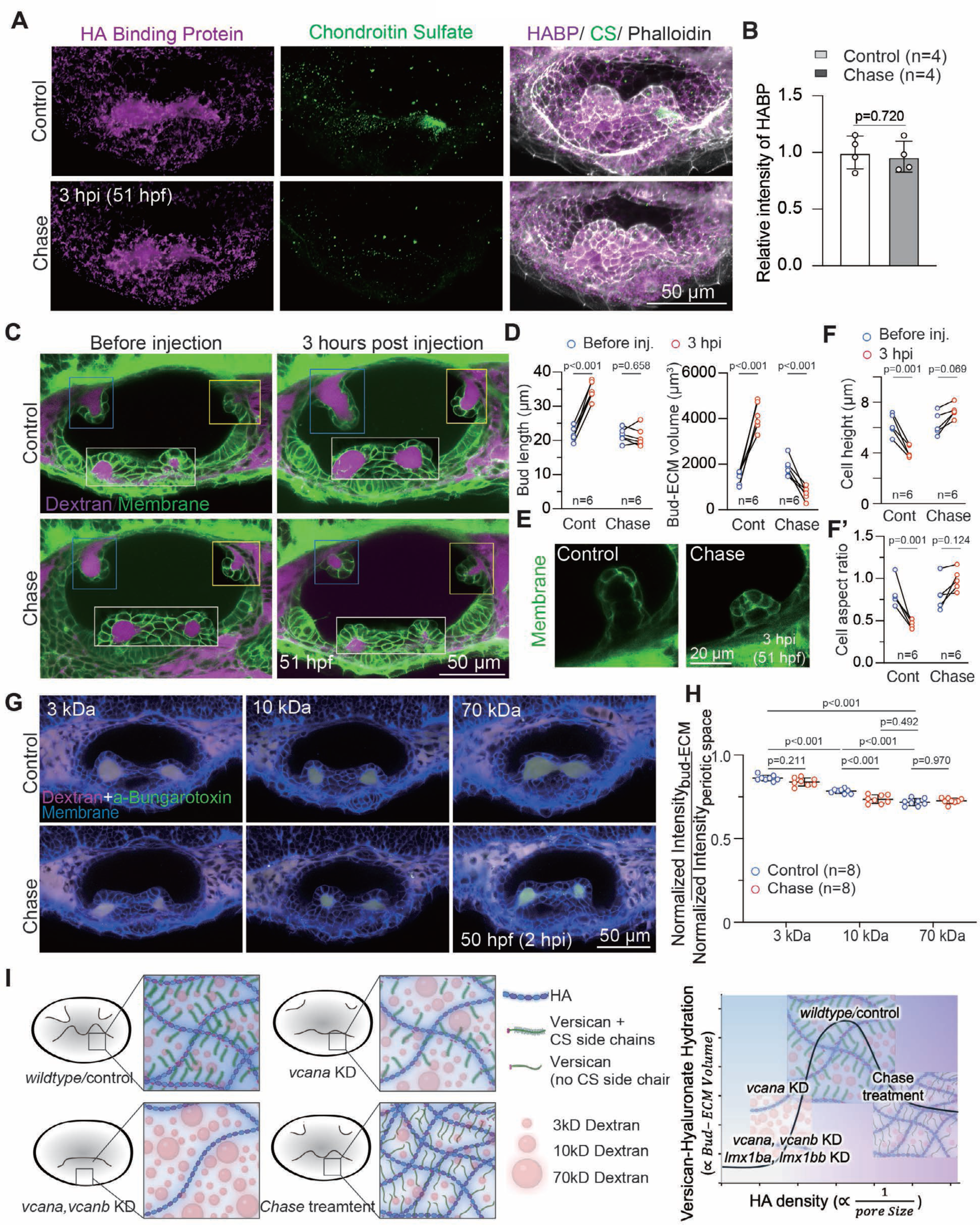
Chondroitin sulfate is required for the hydration of Versican-hyaluronate ECM. **A)** 3D-rendered OVs showing hyaluronic acid (HA), chondroitin sulfate, and F-actin using HA-binding protein, anti-chondroitin sulfate antibody, and phalloidin, respectively, in embryos injected with control buffer or chase (10 U/mL) at 3 hpi. Scale bar, 50 μm. **B)** Quantification of fluorescent intensity of HABP in embryos injected with control buffer of Chondroitinase (Chase) at 3 hours post injection (hpi). 5 μm-thickness maximum intensity projection images of HABP in the lateral bud region were used for the quantification. Data are mean ± SE. “n” denotes the number of embryos measured per condition. p values as labeled (paired two-tailed Student’s t-test). **C)** Composites of 2D sections from membrane-NeonGreen-expressing embryos before injection or at 3hpi of control buffer or Chase. Pre-injection and post-injection images were acquired from the same embryo. Bud-ECM region was labeled with 3 kDa Texas-red Dextran. Anterior, posterior, and lateral buds from different z-depths are framed in blue, yellow, and white boxes, respectively. Scale bar, 50 μm. **D)** Quantification of bud length (left), bud-ECM volume (right) at anterior bud in embryos before injection or at 3hpi of control buffer or Chase. Bud-ECM volume was measured using the bud-ECM region labeled by Texas-red Dextran. Data are mean ± SE. “n” denotes the number of embryos measured per condition. p values as labeled (paired two-tailed Student’s t-test). **E)** Representative 2D sections of anterior bud in membrane-NeonGreen-expressing embryos injected with control buffer or chase at 3 hpi. **F)** Quantification of cell height (F) and cell aspect ratio (F’) at anterior bud in the membrane-NeonGreen-expressing embryos before injection or at 3 hpi of control buffer of chase. Data are mean ± SE. “n” denotes the number of embryos measured per condition. p values as labeled (unpaired two-tailed Student’s t-test). **G)** Composite of 2D sections of OVs showing percolation of dextran from periotic region to into bud-ECM region at 2 hpi. Control buffer or Chase was co-injected with Texas-red dextran (magenta: 3, 10, or 70 kDa) and aBt (green) in embryos expressing membrane-NeonGreen (blue). aBt colocalizes with all three dextran sizes in the periotic region (white). Contrast is the same across embryos. Experiments with 10 kDa dextran were performed on different days than experiments with 3 kDa and 70 kDa dextrans. Scale bar, 50 μm. **H)** Quantification of fluorescent intensities of different sizes of dextran in the bud-ECM region normalized to their intensities in the periotic space. “n” denotes the number of embryos measured per condition. p values as labeled (one-way ANOVA with Tukey’s test).**G)**A schematic model for the regulation of bud-ECM density and hydration regulated by Versican and its CS side chains.

### Versican regulates the density of the bud-ECM via the accumulation of HA

We next addressed the underlying mechanism of Versican function during canal morphogenesis. Versican’s protein core has a conserved G1 domain with an HA-binding site(LeBaron et al., 1992). The non-covalent Versican-HA interaction is often stabilized by link proteins, such as *hapln1a*, which is also expressed in the canal genesis zones of the inner ear(Geng et al., 2013). Versican-HA aggregates form deformable secondary structures, e.g., lattice-like structures with increased viscosity in arterial smooth muscles(Evanko et al., 2001). HA and Versican can exist separately but often co-localize in most ECMs in which they are expressed (Wight et al., 2020). While there is no available zebrafish antibody for the Versican protein core, its CS side chains are known to accumulate in the ECM underneath the canal-genesis zones of the OV (Jones et al., 2022). Using a commercially available binding protein (HABP), HA has also been shown to accumulate locally in this ECM to drive the budding of the overlying epithelium through an osmotic swelling (Jones et al., 2022; Munjal et al., 2021). We hypothesized that one of the key functions of Versican is to regulate the density of HA in the bud-ECM. We measured HA levels in the bud-ECM using the HABP in controls and *vcana*, *vcanb* double knockdown embryos. Strikingly, HABP intensity in the bud-ECM of the OV in double knockdown embryos was reduced by 50% (**Figures 4A-B**). *Vcana* and *vcanb* loss-of-function also caused a loss of CS in the ECM of the bud (**Figure 4A**). These data show that Versican is necessary for HA and CS accumulation in the bud ECM.

The cross-linked aggregates in hyaluronate-rich gel matrices have a small pore size of a few tens of nanometers, indicating low permeability and high density (Chaudhuri et al., 2020; Cowman et al., 2015; Munjal et al., 2021). Given the evidence above that Versican is required for HA and CS accumulation in the bud-ECM, we tested if Versican exhibits modulatory control over the density of the bud-ECM by measuring its permeability. To investigate this quantitatively we used two different sizes of red fluorescent-labeled dextran, 3kDa and 70kDa, with a hydrodynamic radius of 1.3nm and 5nm, respectively. Specifically, we injected the dextrans into the space surrounding the OV (peri-otic space) and measured their percolation in the bud-ECM (**Figure 4C**). We co-injected a far-red labeled alpha-Bungarotoxin (aBt), which freely permeated and equilibrated in the bud-ECM relative to the periotic space, to normalize the intensity of the injected and percolated dextran. Instead of a double knockdown of *vcana* and *vcanb* that causes a dramatic loss of HA and CS (**Figure 4A-B**), the percolation assay was carried out in single *vcana* MO-injected embryos. The 3kDa dextran percolated freely in the bud-ECM of control embryos, while the 70kDa dextran did not (**Figures 4D-E**), as previously reported(Munjal et al., 2021). In contrast, both 3kDa and 70kDa dextrans percolate freely in *vcana* MO-injected embryos (**Figures 4D-E**), showing that the bud-ECM has a larger pore size in the Versican knockdown. We conclude that Versican can modulate the permeability and density of the bud-ECM.

### Chondroitin sulfate is required for the hydration of Versican-hyaluronate ECM

Next, we investigated if the chondroitin sulfate (CS) side chains on Versican play a distinct role in morphogenesis compared to its protein core. Previous studies have reported aberrant cardiac and semicircular canal morphogenesis in *chsy1* MO-injected embryos(Li et al., 2010; Peal et al., 2009), suggesting a separate role for CS. HA drives the budding of the canal-genesis zone through osmotic swelling. CS, like HA, is negatively charged. Therefore, it can contribute to the osmotic swelling and hydration of the Versican-hyaluronate aggregates(Islam and Watanabe, 2020; Maroudas, 1976; Richter et al., 2018). To test this role, we injected Chondroitinase ABC (Chase) to digest CS in the bud-ECM. At 10 U/mL concentration of Chase injected in the peri-otic space, we detected no change in the HABP signal and complete inhibition of CS in the bud-ECM, which showed specificity to CS (**Figures 5A-B**). The extension of buds and, importantly, the bud-ECM volume decreased in these embryos compared to vehicle controls (**Figures 5C-D**). Previous studies have reported that due to the incompressible nature of water, osmotically charged hydrogels can apply hydrostatic pressure on the surrounding tissue(Chaudhuri et al., 2020; Nakamura and Manasek, 1981; Toole, 2001; Vignes et al., 2022). Consistent with this, the columnar cells change their morphology from tall to flat, consistent with stretching during extracellular pressure-driven extension (**Figures 5E-F**) (Munjal et al., 2021). Strikingly, when CS was digested with Chase, cell stretching was reduced, as measured by the cell height and aspect ratio, showing a reduction in the extracellular pressure (**Figures 5E-F**). Together, these data confirm the contribution of CS to the osmotic swelling and pressure buildup in the Versican-hyaluronate ECM.

CS digestion does not perturb HA accumulation and yet caused a 50% reduction in the bud-ECM volume. We therefore hypothesized that lower osmotic swelling would lead to a more compact bud-ECM with lower permeability. We tested this hypothesis using the dextran percolation assay described above, injecting 3kDa, 10kDa, and 70kDa dextrans in the periotic space. The 3kDa and 10kDa dextrans percolated freely in controls, while the 70kDa dextran did not (**Figures 5G-H**). A small but significant reduction in the percolation of 10kDa dextran was observed in Chase-injected embryos, making it comparable to 70kDa percolation in controls (**Figures 5G-H**), indicating a smaller pore size. Notably, the effect of CS digestion resulting in lower dextran percolation is distinct from the effect of *vcana* knockdown resulting in higher dextran percolation (**Figures 4D-E**), consistent with HA accumulation in these conditions (unchanged or lower). Altogether, we conclude that while Versican can modulate the accumulation and density of the bud-ECM through non-covalent interaction with HA via its protein core, the presence of covalently attached CS-side chains contributes to the osmotic swelling and hydration of Versican-hyaluronate rich bud-ECM (**Figure 5I**).

## Discussion

The correct morphogenesis of the semicircular canals is absolutely essential for their physiological function. Recently, we and others discovered a key role for hydrostatic pressure from the hyaluronate-rich ECM in providing the force necessary for bud outgrowth in the non-sensory domain of the zebrafish OV (Jones et al., 2022; Munjal et al., 2021). While the expression of genes involved in the synthesis of HA and HA binding proteins, such as Versican, had been documented previously in this domain, the factors controlling their precise temporal and spatial expression and the relationship of this expression to Notch signaling were not known. Moreover, how the different matrix components co-operate to modulate hydrostatic pressure was also a completely open question since little emphasis has been placed on the role of nonprotein components of the ECM in tissue morphogenesis. Here, the advantages of the zebrafish as a model organism have enabled us to throw light on all these critical questions using a combination of genetic perturbations and high-resolution morphogenetic analysis.

The first key finding of this work is the identification of the transcription factor Lmx1b, encoded by two paralogs, *lmx1ba* and *lmx1bb*, as a positive transcriptional regulator of HA and Versican synthesis during semicircular canal morphogenesis. We also identify a role for Notch signaling in the sensory domain in the patterning of *lmx1b* in the non-sensory domain, akin to the antagonistic interactions between Notch and Lmx1a in embryonic mice and chicken inner ear(Mann et al., 2017). We show that loss of Notch signaling broadened the expression domain of *lmx1bb* while suppressing the expression domain of *lmx1ba.* The differential expression pattern of the two paralogs suggests combinatorial gene regulation that remains to be investigated.

In humans, mutations in LMX1B are associated with a pleiotropic disorder called nail patella syndrome, which is associated with nail and skeletal malformations, glaucoma, and kidney disease (Bennett et al., 1973; Dreyer et al., 1998; Mimiwati et al., 2006). Similarly, mouse models with *Lmx1b* knockout have renal, ocular, and limb abnormalities(Gu and Kania, 2010; Krawchuk and Kania, 2008). Finally, Lmx1b is indirectly involved in digit regeneration in mice(Castilla-Ibeas et al., 2024; Johnson et al., 2022), an ability possessed by many rodents and some primates, including human children. ChIP-seq (chromatin immunoprecipitation followed by sequencing) data from embryonic mouse limbs has identified several genes with *cis*-regulatory modules that bind to Lmx1b(Haro et al., 2017). These include ECM genes such as *Col1a2, Vcan,* and *Has3*. Together, these studies and our findings suggest a broad role of Lmx1b in regulating ECM biosynthesis during the development and possibly regeneration of organs. It remains to be elucidated how Lmx1b specifically activates the expression of glycosaminoglycan and proteoglycan biosynthesis during semicircular canal morphogenesis. Another important question remains whether Lmx1b is sufficient for expressing ECM genes or if other developmental factors are involved.

Versican-hyaluronate ECMs make a transient appearance in several tissues during dynamic morphogenesis events (Wight, 2017). For example, Versican is dynamically expressed during cardiac development, and several cardiovascular defects, and even lethality, are associated with Versican loss or mis-expression(Mjaatvedt et al., 1998; Nandadasa et al., 2014; Yamamura et al., 1997). Similarly, Versican is expressed during eye development(Kwan, 2014), and in humans, a hereditary disorder named Wagner syndrome causes progressive vision loss due to mutations in the *VCAN* gene(Kloeckener-Gruissem et al., 2006). Yet the molecular mechanisms by which Versican is required for healthy organ development and the etiologies of these diseases have remained largely unknown. Our second key finding in this study is that Versican and its CS side chains have two distinct functions during zebrafish canal morphogenesis. The first function is that Versican, via its protein core, regulates the accumulation of HA and, thereby, bud-ECM density. Versican-dependent HA accumulation is indispensable for successful canal morphogenesis as it depends on hyaluronate pressure for tissue remodeling. The second function of Versican is that it tunes the osmotic swelling and hydration of the hyaluronate-rich ECM via its negatively charged sulfated chondroitin chains (**Figure 5I**). The broad implications of this function are discussed below.

The number of sulfate side chains attached to proteoglycans directly impacts the properties of the hyaluronate-rich matrices they interact with (Richter et al., 2018). For example, cartilage resistance to compressive stresses is due to the swelling potential of the large proteoglycan, aggrecan, that has an abundance of chondroitin and keratin sulfate chains and forms secondary structures with hyaluronate and collagen fibrils(Maroudas, 1976; Roughley and Mort, 2014). The conserved CS binding domain in Versican’s protein core is alternatively spliced in most species in a context-dependent manner, leading to isoforms with two, one, or no subdomains for CS attachment(Dours-Zimmermann and Zimmermann, 1994; Paulus et al., 1996). While there is no evidence of alternative splicing in zebrafish, the *vcana* paralog encodes a protein that appears to lack the CS binding domain, while the *vcanb* paralog encodes a protein with both subdomains for CS binding(Derrick and Noel, 2021). We hypothesize that the differential expression of *vcana* and *vcanb* in zebrafish or alternative splicing of Versican in other species is a key mechanism underlying its modular functions. More CS side chains would add osmotic charges to hyaluronate-Versican aggregates, filling up larger volumes through hydration or building up more hydrostatic pressure if confined. Fewer CS side chains would reduce osmotic charges, making more compact ECMs with low permeability to signaling molecules. This hypothesis can be investigated in other organs where only one paralog or spliced isoform of Versican is expressed (such as *vcana* in the developing heart, lens, and tail primordium(Thisse and Thisse, 2008)). Besides the HA and CS binding domains, Versican has a third conserved domain that can interact with other extracellular components, such as fibrillins and integrins(Dongning et al., 2022; Wu et al., 2005). The functions of these interactions in semicircular canal morphogenesis remain to be investigated.

The primary roles of the ECM in tissue morphogenesis have been attributed to its protein components, such as collagens, fibronectins, and laminins, which form stiff fibrous polymer networks that transmit tensile forces. However, increasing evidence, including our findings here, shows that the ECMs are not simple elastic materials. Instead, many physiologically relevant ECMs incorporate GAGs and proteoglycans that can confer on them tunable viscoelastic properties (Chaudhuri et al., 2020; Cowman et al., 2015).

In conclusion, our research has revealed two important new features during a healthy developmental process: a critical role for Versican in building a tunable viscoelastic gel matrix with hyaluronate polymers to drive tissue remodeling, and a role for the transcription factor Lmx1b in promoting the expression of a proteoglycan-rich ECM during canal morphogenesis. These findings have direct implications for diseases where these matrices are impaired and for improving biomaterials used in organoid engineering and regenerative medicine.

## Materials and Methods

### Animals

Zebrafish (*Danio rerio)*–EK and AB wildtype strains were used in this study. Adult fish were kept on a 14-hour light/10-hour dark cycle. Embryos were collected by crossing female and male adults (3-18 months old) and raised in 28⁰.5 °C egg water. In this study, the following transgenic lines were used:

1. *Tg(actb2:membrane-neongreen-neongreen)(Munjal et al., 2021)*,
2. *Tg(*β*Actin:membrane-mCherry)(Xiong et al., 2014)*
3. and *vcana ^pd3001/pd3001^, vcanb ^pd3002/pd3002^* (this study), which we name *memai*, Japanese for “dizzy”(めまい or 目眩)

All experiments with animals were carried out in compliance with internal regulatory review at Duke University School of Medicine.

### Genome editing by CRISPR-Cas9

***1) lmx1b* crispant embryo**

For knockdown of either *lmx1ba* or *lmx1bb*, 3 single guide RNAs (sgRNAs) targeting each gene were pooled together. *lmx1ba* sgRNAs target multiple coding regions of the *lmx1ba* gene. *lmx1bb* sgRNAs target exon 5 in the *lmx1bb* gene. 200 ng/uL of sgRNAs pool targeting either *lmx1ba* or *lmx1b* were injected with 500 ng/ uL of Cas9 protein at the single-cell stage. For the double-knockdown of *lmx1ba* and *lmx1bb*, 6 sgRNAs were pooled together. A working solution containing 33 ng /uL of each sgRNA and 500 ng/uL of Cas9 protein was injected as described above. F0 crispant embryos were screened with inner ear phenotype and T7 endonuclease assay at 50 hpf and then used for confocal imaging with live embryos or HCR-FISH at 50-72 hpf.

The following target sites were used to generate each sgRNA:

1. *lmx1ba* sgRNA1: 5’ CCTCACGCCGCCGCAGATGCCCG 3’
2. *lmx1ba* sgRNA2: 5’ CCGACCCTCTCGACAGATCCGGC 3’
3. *lmx1ba* sgRNA3: 5’ GAGAAAGGCGCTGGCGGTCAGGG 3’
4. *lmx1bb* sgRNA1: 5’ AGGCTCGGGAGGCACCGGGAAGG 3’
5. *lmx1bb* sgRNA2: 5’ GAGAAAGGCTCGGGAGGCACCGG 3’
6. *lmx1bb* sgRNA3: 5’ CCCTGCCGGAAGGTGCGTTCCTC 3’

A T7 endonuclease assay was performed using the following primers:

1. *lmx1ba* F:5’GCTCAGCAGAATGACGCGAT 3’; R: 5’ CGCGTTTAAAGATCTGCGTGTA 3’
2. *lmx1bb* F:5’ CCACGTACCATCCTCACCAC 3’; R: 5’ TGCAATGTCCCAAATTCGCT 3’

Heteroduplex mobility shift assay was performed to test guide activity using the following primers:

3. *lmx1ba* sgRNA1 F:5’ ATGGAGAGCAGCGGATACAG3’; R: 5’ TGATCTTCACACAGGGCTCA 3’
4. *lmx1ba* sgRNA2 F:5’ ACATACACACACCGCTGGAT 3’; R: 5’ TGATCACACCGACTCTGAGG 3’
5. *lmx1ba* sgRNA3 F:5’ GGATGAAGATCTGGATGTGAAGC 3’; R: 5’ GATGGTTCGGGGTCTTTTGG 3’
6. *lmx1bb* sgRNA1 and sgRNA2 F:5’GCGAAGATGAGGAGCTGGAT 3’; R: 5’ ATGGTACGTGGTCTCTTGGG3’
7. *lmx1bb* sgRNA3 F:5’ ACCACACAGCAGAGACGAG 3’; R: 5’ ACAAGAGATGGAGGAAGCTGA3’

***2) vcan* crispant embryo**

For the knockdown of either *vcana* or *vcanb*, 3 sgRNAs were generated and pooled, as mentioned above. *vcana* and *vcanb* sgRNAs target multiple *vcana* and *vcanb* coding regions, respectively. 200 ng/uL of sgRNAs pool targeting *vcana* or *vcanb* were injected with 500 ng/ uL of Cas9 protein at the single-cell stage. For the double-knockdown of *vcana* and *vcanb*, 6 sgRNAs were pooled together, and then 33 ng /uL of each sgRNA and 500 ng/uL of Cas9 protein were injected as described above. F0 crispant embryos were screened with inner ear phenotype at 50 and then used for confocal imaging with live embryos or HCR-FISH at 50-72 hpf.

The following target sites were used to generate each sgRNA:

1. *vcana* sgRNA1: 5’ GTGGTGGTCTGCTGGGAGTACGG 3’
2. *vcana* sgRNA2: 5’ TGGGATCTGGGAAGCCTGTTTGG 3’
3. *vcana* sgRNA3: 5’ GTGTTAGGTCGTATGGCAAACGG 3’
4. *vcanb* sgRNA1: 5’ GGGTAAAGGGTAGCCCGTCCGG 3’
5. *vcanb* sgRNA2: 5’ GTCGCTGTGGTGGCCCAGAACGG3’
6. *vcanb* sgRNA3: 5’ GTGTGAAGTTCAGACTGTAGCGG 3’

The following primers were used for heteroduplex mobility shift assay to test guide activity:

1. *vcana* sgRNA1 F:5’ GGCAGTGCACGCCATC 3’; R: 5’ CCTCATACTGGGATCTGGGAAG 3’
2. *vcana* sgRNA2 F:5’ GCTTTCTGATGGCAGTGCAC 3’; R: 5’ CCTTGGAAACAGTATGCGCC 3’
3. *vcana* sgRNA3 F:5’ TGCCTGGAAAACCTGGTGTT 3’; R: 5’ AGAACTGTTGGCCTTGTCAAC 3’
4. *vcanb* sgRNA1 F:5’ ATACCCGGTCTCCGTTCC 3’; R: 5’ CACCTCTGAAACAGTAGGCTCC 3’
5. *vcanb* sgRNA2 F:5 TGATCACCTGCGCATCAAGT 3’; R: 5’ GATGGTCAGAGAAGCGTCCC 3’
6. *vcanb* sgRNA3 F:5’ TTTTGCTGGTGTCTGCAGGA 3’; R: 5’ TAAAGCTGTTCAGGGGTGGC 3’

***3) tyrosinase* crispant embryo**

As a control embryo, 200 ng /uL of a sgRNA targeting *tyrosinase* gene(Sorlien et al., 2018) was injected with 500 ng/uL of Cas9 protein. Cas9-induced indel formation at the *tyrosinase* gene results in loss of pigmentation. The *tyrosinase* crispant embryos were screened with loss of pigmentation at 50 hpf and used for confocal imaging with live embryos or HCR FISH at 50-72 hpf. The following target sites were used to generate a sgRNA:

*tyrosinase* sgRNA: 5’ GGACTGGAGGACTTCTGGGG 3’

### memai mutant (vcana ^pd3001/pd3001^, vcanb ^pd3002/pd3002^) generation

The mutations in the HA-binding domain of both *vcana* and *vcanb* were generated using CRISPR-Cas9. *vcana* sgRNA1 and *vcanb* sgRNA1 were pooled together, and then 200 ng/ uL of the sgRNA pool was injected with 500 ng/uL of Cas9 protein at the single-cell stage. The *vcana* mutant allele resulted in 7 bp intragenic deletion in exon 4, generating a frameshift that induces a premature stop codon after amino acid (aa) 305. The *vcanb* mutant allele resulted in 7 bp intragenic deletion in exon 6, generating a frameshift that induces a premature stop codon after aa 335. The *memai* adults are maintained as *vcana^+/pd3001^*; *vcanb ^pd3002/pd3002^* genotypes. Double knockout embryos of *vcana* and *vcanb* were screened from *vcana ^+/pd3001^; vcanb ^pd3002/pd3002^* parents through characteristic small buds in the otic vesicle and pericardial edema phenotype.

### RNA extraction and reverse transcription PCR (RT-PCR)

30 embryos injected with sgRNAs/Cas9 were collected at 53 hpf. Total RNA was extracted using TRizol reagent (Thermo Fisher Scientific) and Direct-zol RNA MiniPrep kit (Zymo Research) according to the manufacturer’s protocol. Reverse transcription cDNA was synthesized using Superscript Ⅳ reverse transcriptase (Invitrogen). PCR was performed with Phusion Flash High-Fedelity PCR Master Mix (Thermo Fisher Scientific) to assess the gene knockdown in crispant embryos. The following primers were used for PCR:

1. *lmx1ba:* F:5’ ATGTGATGTCCAGCCGGATG 3’; R: 5’ CCGATGCGCTCATGAAACAG 3’
2. *lmx1bb:* F:5’ GAGACAGGTCTGAGCGTACG 3’; R: 5’ TAGCCGTTCTCCAGTGCAAC 3’
3. *vcana:* F:5’ TGTCACACAAACCCTTGTCGT 3’; R: 5’ CCCCTGGAGACGACATTCAC 3’
4. *vcanb:* F:5’ CCACCCCTAAAGCCACAAGT 3’; R: 5’ AAGCATGTTCCGCCGTTTTC 3’
5. *actb2:* F:5’ AGAGCTACGGAGCTGCCTGAC 3’; R: 5’ TACCGCAAGATTCCATACCC 3’

### Morpholino injection

Standard control morpholino oligonucleotides (MO) and previously characterized MOs targeting *vcana*: 5’-CTGAAACACCCATGGGAGTGGACAT-3’(Chen et al., 2012; Muller-Deile et al., 2016), *vcanb*: 5’-TTCACGTCCAACAACATCATCTCAA-3’(Kang et al., 2004), *jag1b*: AATAGTCTTTCTTACGGGAGTGGC(Obholzer et al., 2012), and *p53* were purchased from Gene Tools. 7 ng of either *vcana* MO, *vcanb* MO, or control MO was injected with 3.5ng of *p53* MO(Gerety and Wilkinson, 2011) into embryos at the single-cell stage. For the double knockdown of *vcana* and *vcanb*, a mixture containing 3.5 ng of each MO was injected with 3.5 ng of *p53* MO into embryos at the single-cell stage. 3 ng of *jag1b* MO was injected with 1.75 ng of *p53* MO into embryos at the single-cell stage. MO-injected embryos were screened with inner ear phenotype at 50 hpf and used for confocal imaging with live embryos and immunostaining at 50-72hpf.

### Immunohistochemistry

The dechorionated embryos at various stages were fixed with 4%PFA/PBS, followed by the permeabilization with pre-chilled acetone as previously described (Munjal et al., Cell, 2021). Embryos were blocked with 5% BSA in 1X PBS containing 0.1% Tween-20 (PBT) at RT for 1 hour and then incubated with primary antibody solution containing anti-chondroitin sulfate antibody (Millipore, monoclonal, mouse, 1:100) and biotinylated-Hyaluronan Binding Protein (Millipore, 1:50) at 4°C, overnight. Embryos were washed with 1X PBS for 15 mins, thrice. Embryos were incubated with a secondary antibody solution containing fluorescent-labeled anti-mouse antibody (Thermo Fisher, 1:200), Streptavidin (Thermo Fisher, 1:500), and Phalloidin (Thermo Fisher, 1:100) at 4°C overnight. Embryos were then washed with PBS for 15 mins, thrice, and mounted for confocal imaging or stored at 4°C.

### Chondroitinase treatment

The dechorionated embryos at 48 hpf were soaked in 1X tricaine and immobilized in a 1.5% canyon mount filled with egg water. Chondroitinase (Chase from Sigma Aldrich) was dissolved in a 0.01% BSA solution and stored at 50 units/mL at -20°C. The injection solution was made with a buffer containing 50 mM Tris-Hcl, 60 mM sodium acetate, 0.02% BSA, and 0.5% Phenol Red. Chase (10 units/mL) was injected into the periotic space (anterior and posterior to the OV) of embryos at 48 hpf. The embryos were then fixed with 4%PFA/PBS at 3 hours post-injection, followed by immunostaining. To measure bud length and bud-luminal volume, 3 kDa Texas-red Dextran (Thermo Fisher, 500 μM) was injected into the periotic space of embryos at 48 hpf, followed by confocal imaging of OVs. Chase (10 units/mL) was then injected with 3kDa of Texas-red Dextran into periotic space, and OVs were captured again by confocal imaging at 3 hours post-injection.

### Confocal imaging

Confocal images were performed with Zeiss LSM 980 using a C-Apochromat 40X 1.2 NA objective for all fluorescent data. For live imaging, dechorionated embryos were anesthetized in 1X tricaine and then mounted dorsolaterally using a canyon mount cast in 1.5% agarose dissolved in egg water, covered with a cover slip, as previously described (Munjal et al. 2021).

### Multiplex *in situ* hybridization chain reaction (HCR)

HCR probes of each gene were obtained from IDT (oligoPools). Hybridization buffer, amplification buffer, wash buffer, and amplifiers were obtained from Molecular instruments. HCR FISH experiments were performed as previously described(Munjal et al., 2021). Dechorionated embryos at various stages were fixed with 4% PFA/PBS and permeabilized with prechilled acetone. The embryos were then incubated with the hybridization buffer at 37°C for 30 min, followed by the incubation with hybridization solution containing 1 pg of probes overnight at 37°C. Fluorescent signals were generated and amplified by probes binding with fluorescent HCR amplifiers in an amplification buffer overnight at RT. Embryos were washed with 0.1% Tween-20 in 5X SSCT (four times) and mounted for confocal imaging or stored at 4°C.

### Dextran percolation assay

Dechorionated embryos at 50 hpf were soaked in 1X tricaine and immobilized in a 1.5% canyon mount cast filled with egg water. Texas-red Dextran (3 kDa and 70 kDa from Thermo Fisher) and AF647 conjugated α-bungarotoxin protein (aBt, Thermo Fisher) were dissolved in 1X PBS and stored at 10 mM at -20°C and 1 mM at -80°C, respectively. The injection solution was made with 1X PBS/0.5% Phenol Red. Texas-red Dextran (either 3 kDa or 70 kDa, 500 μM) was co-injected with AF647 conjugated aBt (Thermo Fisher, 500 μM) into the periotic space (anterior and posterior to the OV). Embryos were imaged at 2 hours post-injection. To measure the Dextran percolation in embryos treated with Chase, Texas-red Dextran (either 3 kDa, 10 kDa, or 70 kDa, 500 μM) and AF647 conjugated aBt (Thermo Fisher, 500 μM) were co-injected with Chase (10 U/mL) into periotic space, followed by imaging of OVs at 2 hours post-injection.

#### Quantification and statistical analysis

##### 1) Bud length measurements

Z-projections covering the middle section of the bud (5 μm z-depth) were used for the measurement. Bud lengths were measured manually using the “straight-line” tool in Fiji(Schindelin et al., 2012). Where budding had not started or in perturbations where budding was affected, bud lengths correspond to average cell height at the anterior or posterior region.

##### 2) Volume measurements

The measurement of basal luminal volume under the buds was performed using ITK-SNAP as previously described(Munjal et al., 2021; Yushkevich et al., 2006). The anterior bud was segmented manually in each slice of the 3D images of the otic vesicle based on basal luminal staining with 3 kDa Texas-red dextran. The number of voxels that belong to the mesh was converted to actual volume using the known voxel resolution.

##### 3) Measurement of the number of embryos with aberrant morphogenesis

Number of embryos injected with sgRNA/Cas9 pools with visible aberrant morphologies by the total number of embryos measured at 72 hpf.

##### 4) Intensity analysis of multiplex in situ HCR

Fluorescent intensity measurements of the HCR probes were performed by Fiji using maximum intensity z-projections covering the entire anterior or posterior bud at various stages. To create a histogram plot of fluorescence intensity, a region of interest (ROI) was manually drawn along the anterior budding region using the “segmented line” tool (20 pixels wide), with 20 microns on either side of the non-budding region surrounding the budding cells. The “plot profile” tool was then used to obtain an average intensity trace across the line. For measuring the mean intensity of HCR probes in anterior buds, ROI was manually drawn along the outer cells of the bud with a segmented line” tool (20 pixels wide).

##### 5) Intensity analysis of HABP staining

Fluorescent intensity measurements of HABP staining in *tyrosinase* and *vcan* crispant embryos were made in Fiji using maximum intensity z-projections covering both lateral buds at 50 hpf. Using the “Rectangle” tool,3 ROIs were drawn manually within the lateral bud, and the mean intensity was calculated.

##### 6) Dextran percolation

Percolation of different dextran sizes was measured in Fiji using maximum intensity z-projections covering all lateral bud-ECM at 52 hpf. Respectably, 3 ROIs in the bud ECM and the periotic space were drawn manually. The mean intensity in the red channel (Texas-red Dextran) was normalized to the mean intensity in the far-red channel (aBt). Percolation was measured as a ratio of the normalized intensity in the bud-ECM to the normalized intensity in the periotic space.

##### 7) Statistical analysis

Average values are calculated from ‘n,’ where ‘n’ represents the number of buds, bud-ECM, and cells (as labeled in each figure). Plots show individual data points, mean values, and error bars (standard deviation or standard error of the mean, as labeled). Before performing parametric statistical tests, the Kolmogorov-Smirnov and Shapiro-Wilk tests were used to test if the dataset in each experiment was from a normal distribution. To compare the difference between the two groups, unpaired or paired two-tailed Student’s t-tests were performed. For multiple comparisons, one-way ANOVA with Tukey’s test was performed. p values less than 0.05 were considered to be significant. All statistical tests and plots were made in GraphPad Prism 10.

## Acknowledgments

We thank Kira Heikes, Nadia Eliora, and all other members of the Principles of Tissue Morphogenesis (Munjal) Lab for their feedback and support. We thank Brigid Hogan and Michel Bagnat for their critical comments on the manuscript. We thank members of the Poss and Bagnat labs for valuable technical input during this project. Biorender was used to make illustrations in Figures 2A, 5I, and Supp Figure 5A. This work was supported by R00HD098918 and DP2115157 to AM. YM is supported by JSPS Overseas Research Fellowship and Duke Regeneration Centre Career Advancement Award.

## Data availability

Information and requests for reagents, resources, and data should be addressed to the corresponding author, Akankshi Munjal. All fish lines generated in this study are available on request without restriction.

## Supplementary Figure Legends

**Figure S1.**
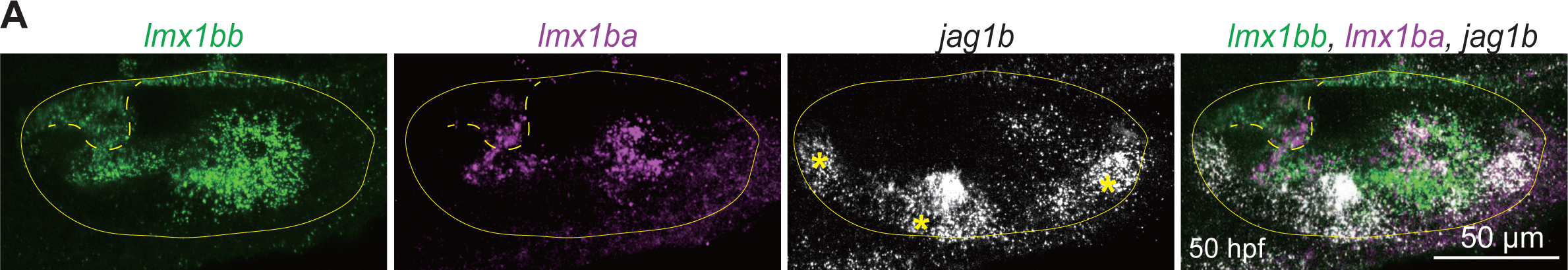
(Related to Figure 1): *Lmx1ba* and *lmx1bb* are expressed in the non-sensory region of the OV, while *jag1b* is expressed in the sensory region of the OV. **A)** Maximum intensity projections of OVs in wildtype embryos at 50 hpf with multiplex *in situ* probes against *lmx1bb* (green), *lmx1ba* (magenta), and *jag1b* (white). The z-volume is set differently to capture buds and cristae. Yellow asterisks mark cristae. The yellow line and the dashed line indicate OV or bud morphology, respectively. Scale bar, 50 μm.

**Figure S2.**
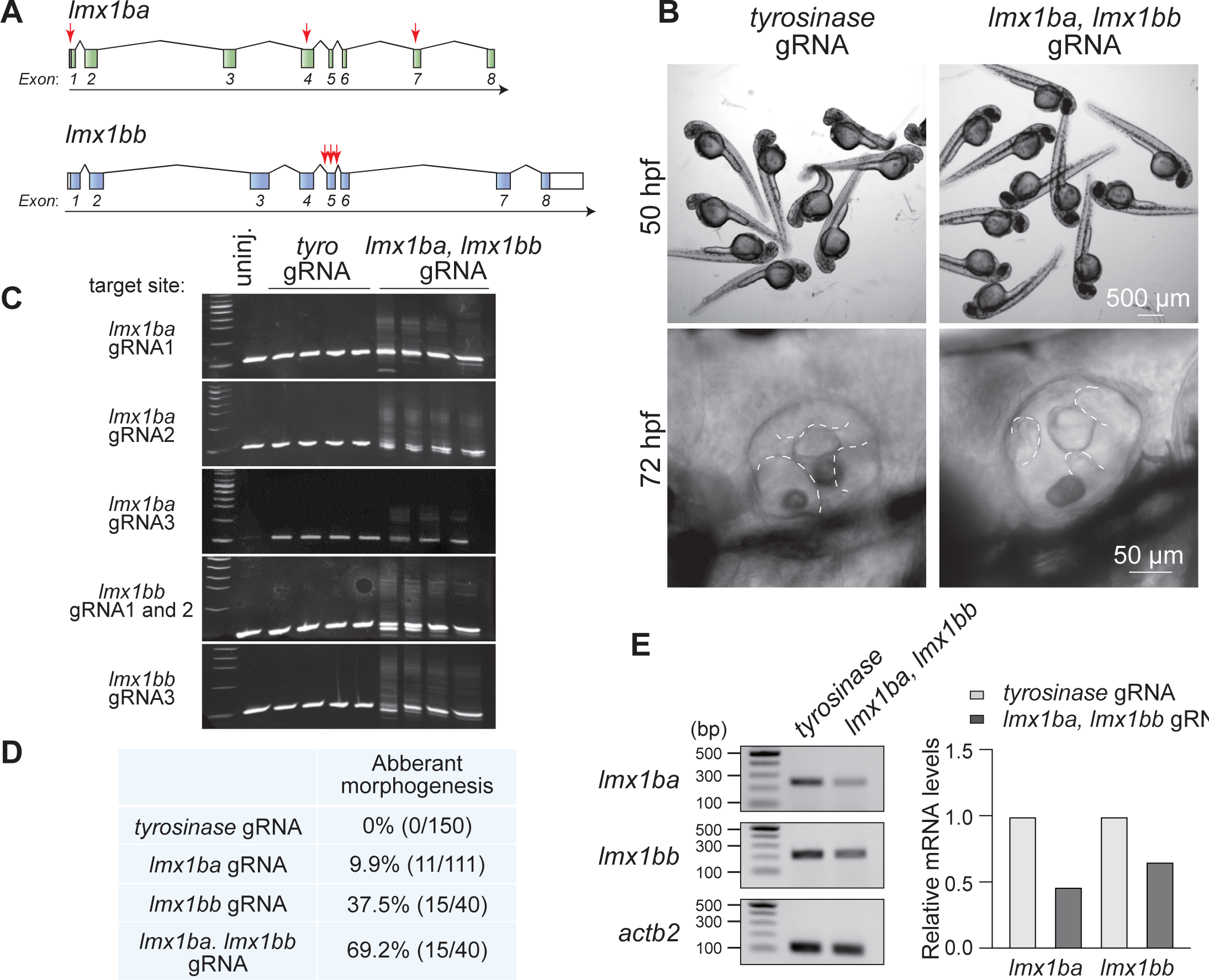
(Related to Figures 1 and 2). CRISPR-Cas9 mediated knockdown of *lmx1ba* and *lmx1bb*, and its effect on HA and Versican synthesis genes. **A)** Target sites for gRNAs at the *lmx1ba* or *lmx1bb* gene locus. Three gRNAs for each gene were injected together with Cas9 protein into the one-cell stage of embryos. Arrows indicate gRNA target sites within exons. **B)** Bright-field images of embryos injected with tyrosinase or *lmx1b*-double gRNA at 50 or 72 hpf. The dashed lines in the magnified images indicate the outlines of pillars or buds in the OV. Scale bar, 50 or 500 μm. **C)** Heteroduplex mobility assay using genomic DNA extracted from embryos injected with t*yrosinase* or *lmx1b*-double gRNA at 72 hpf. Heteroduplexes were observed only in *lmx1b*-double gRNA-injected embryos. Primer pairs targeting the *lmx1ba* or *lmx1bb* coding sequence were used for PCR. **D)** Table showing the number of embryos over the total number measured with aberrant SCC morphogenesis in embryos injected with *tyrosinase*, *lmx1ba*, *lmx1bb*, or *lmx1b*-double gRNA at 72 hpf. **E)** RT-PCR validation of *lmx1ba* and *lmx1bb* knockdown in *lmx1b*-double gRNA-injected embryos. 30 embryos injected with *tyrosinase* or *lmx1b*-double gRNA at 53 hpf were used for total RNA extraction, followed by cDNA synthesis. The mRNA levels of *lmx1ba* and *lmx1bb* were quantified using the band intensity of *lmx1b* genes normalized to *actb2* gene. Relative mRNA expressions are expressed as a ratio relative to gene expressions in *tyrosinase* gRNA-injected embryos. Data are from one experiment.

**Figure S3.**
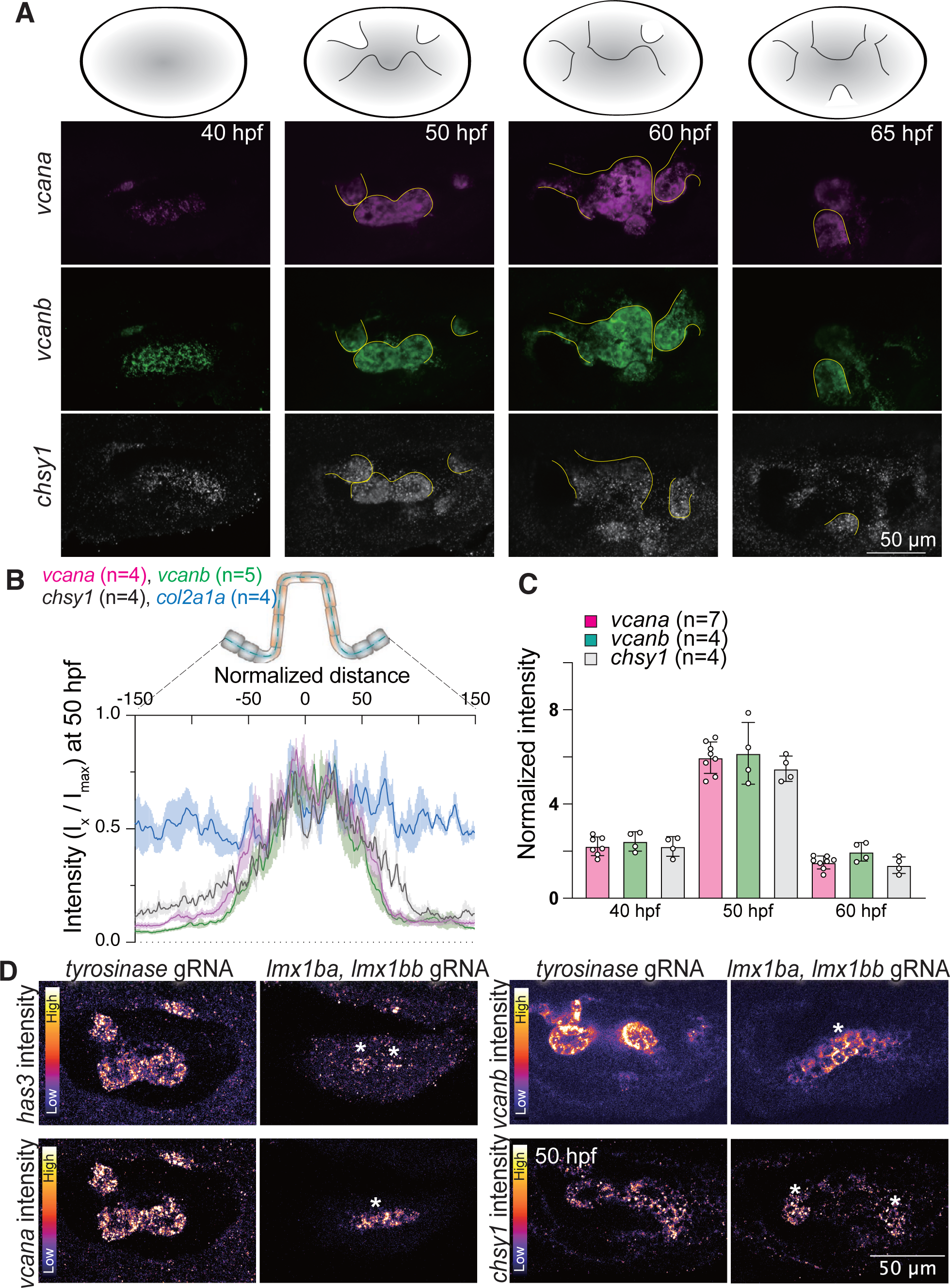
(Related to Figure 2). Expression and patterning of genes encoding Versican, *vcana* and *vcanb,* in OV during canal formation. **A)** 3D-rendered OVs at select time points stained with multiplex *in situ* probes against *vcana* (magenta), *vcanb* (green), and *chsy1* (white). Illustrations show typical morphological features of the OVs at each stage during semicircular canal morphogenesis. The 3D-rendered image stained with the *chsy1* probe is of a different embryo than those stained with the *vcana* and *vcanb* probes. Yellow lines indicate buds or pillar structures in OVs at each time point. Scale bar, 50 μm. **B)** Quantification of fluorescent intensities of various genes across the illustrated region of interest (ROI) in the anterior bud at 50 hpf. Data are mean ± SE. **C)** Quantification of fluorescent probe intensity of *vcana*, *vcanb*, and *chsy1* in the anterior bud at each stage. Data are mean ± SE. “n” denotes the number of buds from individual embryos measured per condition. **D)** Effect of *lmx1b*-double knockdown on ECM-associated gene expression in OV. Maximum intensity projections of *has3*, *vcana, vcanb*, and *chsy1* probes. The z-volume of each condition is individually set to capture lateral buds. Heatmaps represent fluorescent intensity with the same contrast for each probe of embryos injected with *tyrosinase* or *lmx1b*-double gRNA. Scale bar, 50 μm.

**Figure S4.**
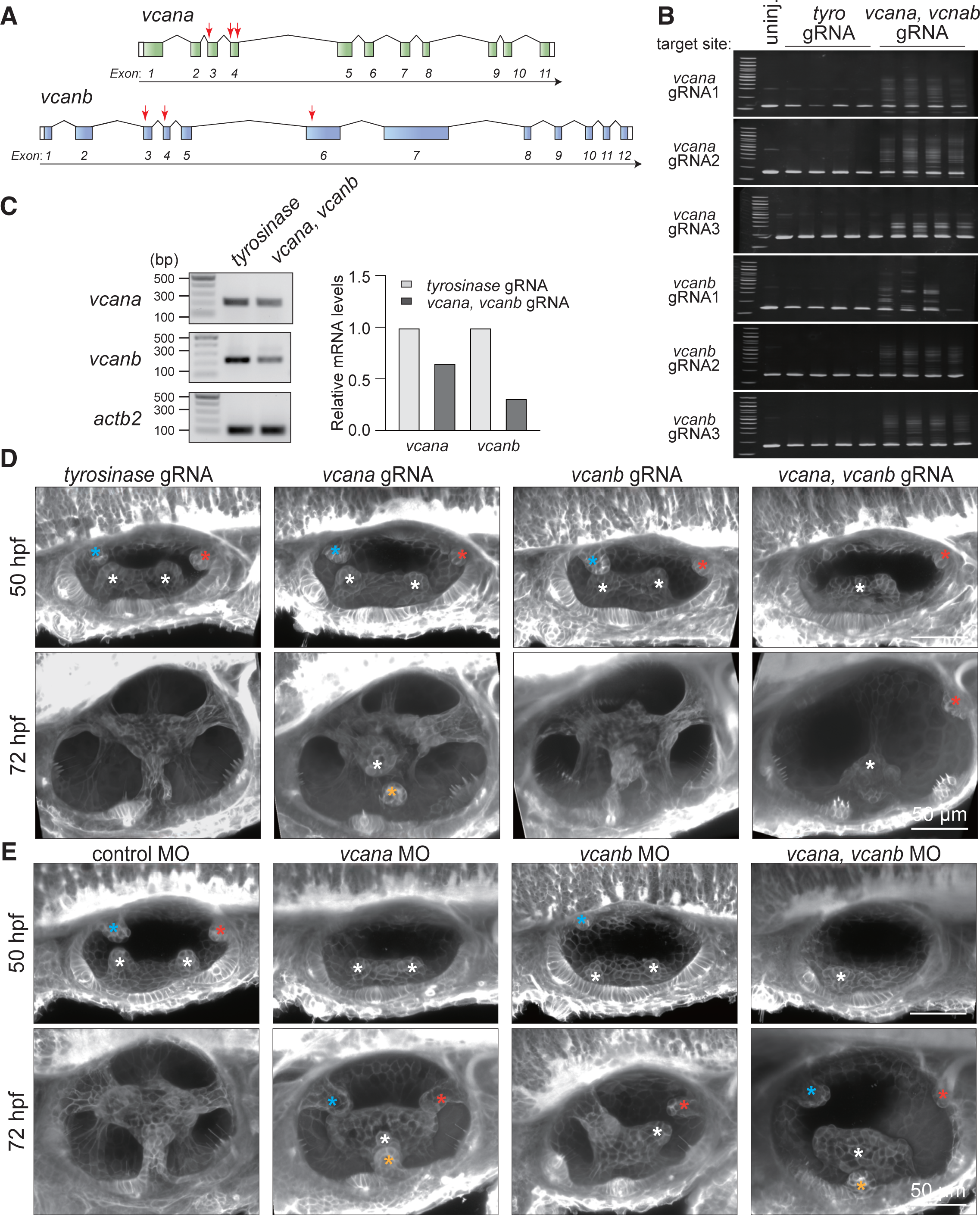
(related to Figure 3): CRISPR-Cas9 and morpholino-mediated knockdown of *vcana* and *vcanb*. **A)** Target sites for gRNAs at the *vcana* or *vcanb* gene locus. Three gRNAs for each gene were injected together with Cas9 protein into the one-cell stage of embryos. Arrows indicate gRNA target sites within exons. **B)** Heteroduplex mobility assay using genomic DNA extracted from embryos injected with *tyrosinase* or *versican*-double gRNA at 72 hpf. Heteroduplexes were observed only in *versican*-double gRNA-injected. Primer pairs targeting the *vcana* or *vcanb* coding sequence were used for PCR. **C)** RT-PCR validation of *vcana* and *vcanb* knockdown in *versican*-double gRNA-injected embryos. 30 embryos of *tyrosinase* or *versican*-double crispant at 53 hpf were used for total RNA extraction, followed by cDNA synthesis. The mRNA levels of *vcana* and *vcanb* were quantified using the band intensity of *versican* genes normalized by the *actb2* gene. Relative mRNA expressions are expressed as a ratio relative to gene expressions in *tyrosinase* gRNA-injected embryos. Data are from one experiment. **D-E)** 3D-rendered OVs at 50 or 72 hpf from membrane-NeonGreen-expressing embryos injected with gRNA (**D**) or MO (**E**). gRNAs or MOs targeting *vcana* or *vcanb* were separately injected into embryos at the one-cell stage to induce *vcana*- or *vcanb*-single knockdown. gRNAs or MO targeting *vcana* or *vcanb* were co-injected at the one-cell stage to induce both *vcana* and *vcanb* knockdown. Anterior, posterior, lateral, and ventral buds are marked by blue, red, white, and yellow asterisks, respectively. Scale bar, 50 μm.

**Figure S5.**
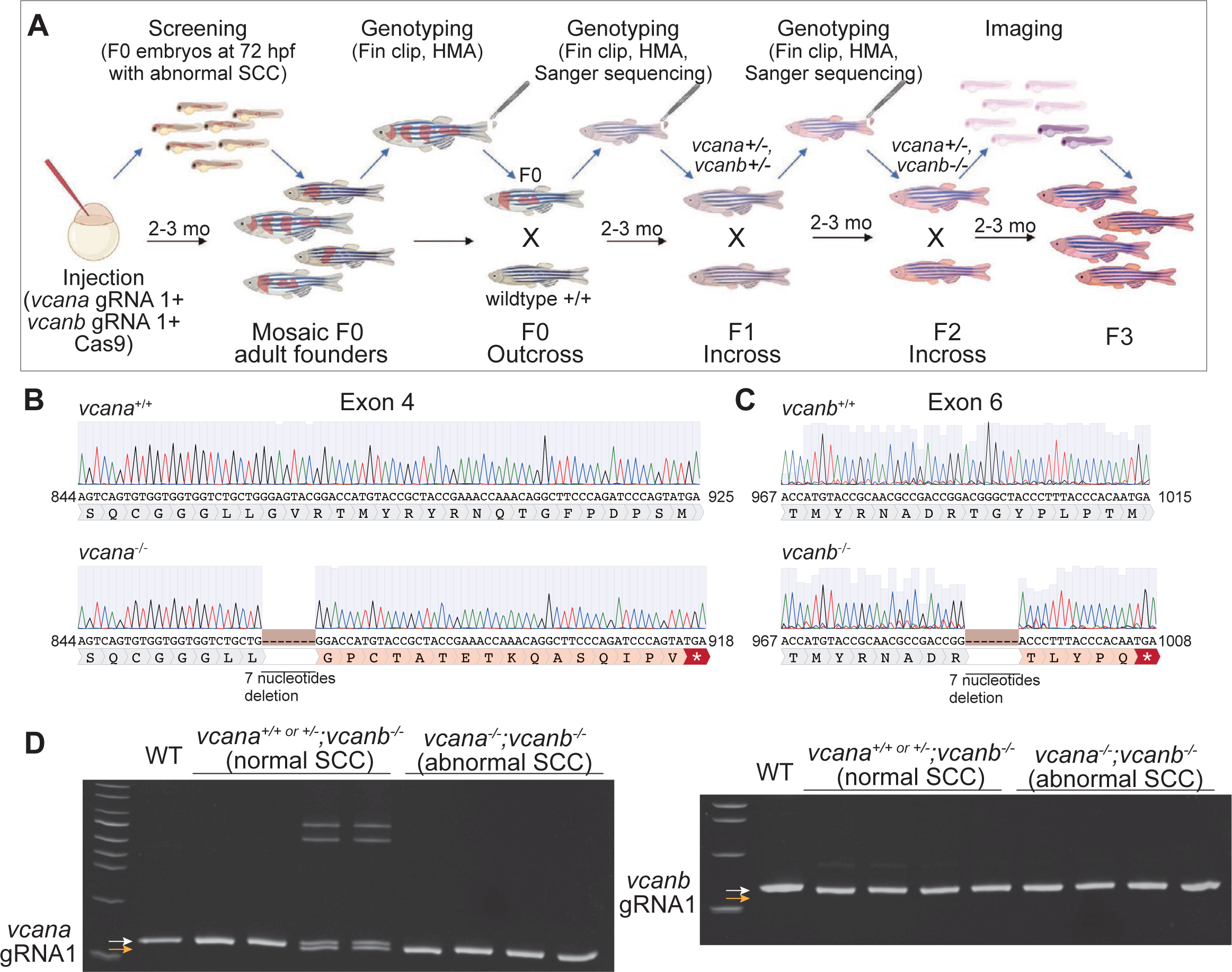
(related to Figure 3): *Memai (vcana^-/-^,vcanb^-/-^)* mutant generation. A) Workflow for screening and validating a stable double mutant of *vcana* and *vcanb*. B) Chromatogram of *vcana ^+/+^* and *vcana^-/-^* sequencing. C) Chromatogram of *vcanb ^+/+^* and *vcanb^-/-^* sequencing. D) Heteroduplex mobility assay using genomic DNA extracted from WT embryos or F_3_ versican mutant embryos obtained from F_2_ vcana^-/+^; *vcanb^-/-^*parents. White and yellow arrows indicate WT or mutant homozygous allele. Primer pairs targeting the *vcana* or *vcanb* coding sequence were used for PCR.

